# *adductomicsR*: A package for detection and quantification of protein adducts from mass spectra of tryptic digests

**DOI:** 10.1101/463331

**Authors:** Josie Hayes, William M. B. Edmands, Yukiko Yano, Hasmik Grigoryan, Courtney Schiffman, Sandrine Dudoit, Stephen M. Rappaport

## Abstract

**Summary:** Liquid chromatography-high resolution mass spectrometry (LC-HRMS) has been used to establish a method, referred to as ‘adductomics’, for characterisation of putative protein adducts at selected loci in human serum albumin (HSA). Applications of this method have been limited by the lack of software for untargeted analysis of modified peptides in protein digests. Here we present *adductomicsR*, an open-source R package for processing LC-HRMS data from analysis of adducted HSA peptides. The software interrogates mass spectra to correct for retention-time drift, and to discover and quantify putative adducts along with those for a housekeeping peptide and internal standard.

**Availability and implementation:** *adductomicsR* is written in R and publicly available at https://github.com/JosieLHayes/adductomicsR, which includes a vignette with example data.

**Supplementary information:** mzXML files for the vignette and test dataset are available in an associated data package adductData (https://github.com/JosieLHayes/adductData).

**Issue Section:** APPLICATIONS NOTE

## 1. INTRODUCTION

Liquid chromatography-high-resolution mass spectrometry (LC-HRMS) can be used to characterise adducts in tryptic protein digests. We previously developed a pipeline for analysing all modifications to Cys34 of the third largest tryptic peptide (T3) of human serum albumin (HSA) (Grigoryan *et al.*, 2016, Lu et al., 2017, Liu, S. et al., 2018). Since manual peak picking and quantitation are labour intensive, we present here an open-source R package (*adductomicsR*) to locate and quantify adducts of the T3 peptide from nano-LC-HRMS data. Although we focus on T3-peptide adducts from HSA, the package is structured to accommodate model spectra from other user-defined peptides and proteins of interest.

## 2. WORKFLOW

*adductomicsR* interrogates tandem mass spectra to find modifications to a user-defined peptide (e.g. the T3 peptide), based on signature ions from a model spectrum that include both fixed (adduct-independent) and variable (adduct-dependent) mass-to-charge ratios (*m/z*). A chemically stable peptide from the same protein, referred to as a housekeeping peptide (HKP), is quantitated to normalise adduct levels for HSA content and digestion efficiency. Peaks corresponding to an internal standard are also measured to adjust for retention-time (RT) drift. For analysis of HSA T3 modifications, the internal standard is an isotopically labelled T3 peptide that has been modified at Cys34 with iodoacetamide and the HKP is a chemically neutral peptide adjacent to T3 (^42^LVNEVTEFAK^51^) (Grigoryan *et al.*, 2016).

An overview of the *adductomicsR* workflow is shown in Figure 1, which addresses each of the following steps in the pipeline.

**Fig. 1.**
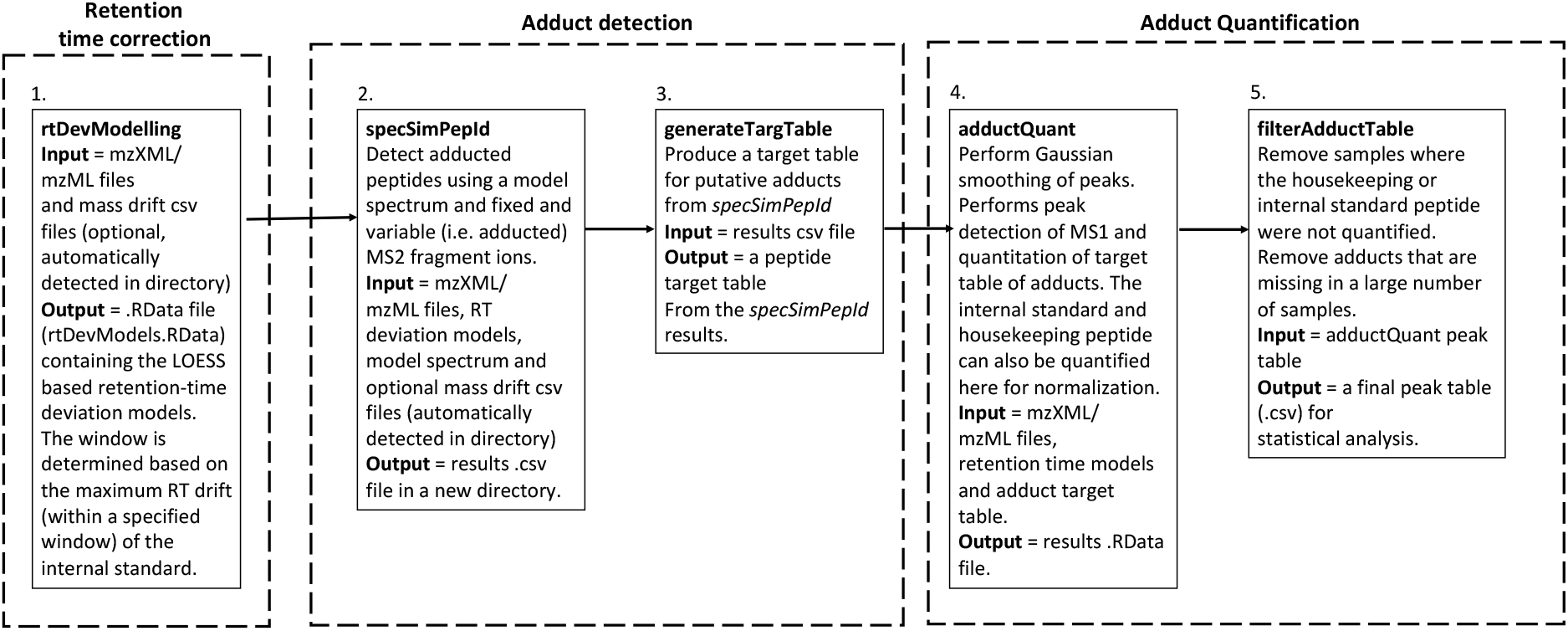
Overview of the *adductomicsR* liquid chromatography-mass spectrometry workflow

### Correction of retention-time drift

Using mzXML as input, *rtDevModelling* corrects for RT drift by determining the RT deviation of known peaks of the internal standard peptide using LOESS models.

### Adduct detection

Peptide spectra are pinpointed by the presence of signature ions from a model spectrum, provided by the user as a csv file, with the *specSimPepid* tool. The relative intensities of fragment ions from each scan are clustered into adduct groups with user-specified RT and monoisotopic mass (MIM) cut points. Colour-coded plots of adduct groups, based on MIM and RT, are produced to assess the validity of the grouping window. Following adduct detection, schematic plots of tandem mass spectra from *specSimPepId* should be visually inspected and the identities of adducts should be manually verified prior to quantification.

### Adduct quantification

*adductQuant* performs peak picking and peak area integration of a target table of putative adducts. The target table can either be defined by the user or generated from LC-HRMS data using *generateTargTable.* Inputs to *adductQuant* are mzXML files plus the RData object, and the output is a .csv file with peak areas of putative adducts. The peak-area ratio of each modified peptide relative to the HKP can be used as a linear measure of the adduct concentration (Grigoryan *et al.*, 2016). A filter step is recommended to remove samples with anomalous abundances of the HKP and/or internal standard, and to remove adducts based on percentages of samples with missing values.

## 3. CONCLUSIONS AND LIMITATIONS

*adductomicsR* is the first open-source package that uses a model spectrum for a peptide to obtain accurate masses of putative adducts, quantitates adduct levels and provides plots of mass spectra to allow the user to assess validity of each putative adduct. The *adductomicsR* package has been tested on Windows and iOS with data files from the T3 peptide of HSA of plasma, serum and dried blood spots from an LTQ Orbitrap XL and an Orbitrap Elite HRMS coupled with a Dionex Ultimate^®^ 3000 nanoflow LC system via a Flex Ion nano-electrospray-ionization source (Thermo Fisher Scientific, Waltham, MA). The method should be readily extrapolated to other mass spectrometers that employ ion traps for MS2.

## Acknowledgements

### Funding

This research was supported by the U.S. National Institutes of Health, via grant R33CA191159 from the National Cancer Institute and grants P50ES018172 and P42ES004705 from the National Institute of Environmental Health Sciences, and by the U.S. Environmental Protection Agency through grant RD83451101.

